# Uncovering Dynamic Neural Information Flow with Continuous-Time Weighted Dynamic Bayesian Networks

**DOI:** 10.64898/2026.01.22.701045

**Authors:** Alec G. Sheffield, Sachira Denagamage, Mitchell P Morton, Anirvan S. Nandy, Monika P. Jadi

## Abstract

Understanding how information dynamically flows within neural systems is a crucial problem in neuroscience. Traditional approaches often assume stationary or quasi-stationary functional networks, which fail to capture the time-varying dynamics of interactions among neural variables. To address this limitation, we introduce Continuous-Time weighted Dynamic Bayesian Networks (CTwDBN), a non-stationary graphical modeling framework for uncovering smoothly time-varying conditional dependencies. Validation on synthetic datasets demonstrated that CTwDBN reliably recovers the structure and dynamics of ground-truth information flow. Application to electrophysiological recordings during a guided saccade task revealed temporal fluctuations in conditional dependencies in the cortical network that persisted an order of magnitude longer than the receptive field dynamics. In the resting-state cortex, CTwDBN revealed persistent fluctuations within a low-dimensional dependency space reflecting canonical anatomical motifs. These results highlight CTwDBN as a versatile analytical framework for capturing dynamic information flow in neural systems with broad applicability to complex biological and artificial systems.

## INTRODUCTION

Information flow in the brain is not static; instead, patterns of interactions between neural populations dynamically reconfigure across different behavioral contexts and internal brain states^1–3^. Inferring this dynamic information flows from the observed neural activity is a central challenge in neuroscience. Traditional analyses that assume stationarity – a fixed network structure that does not change over time – have been previously used to capture these dynamics using well-defined “states” as identified by behavior (resting state^4^, sleep^5^, attention^6^), task (working memory^7^), or the state of the sensory environment (stimulus present/absent^8^). However, these are insufficient to capture the continuous dynamics of behavior, such as fluctuations in attention^9,10^, impulsivity^11^, and those shaped by fluctuations in internal brain states, such as arousal^12,13^. This limitation highlights the need for computational methods that can infer the evolving structure of neural dependencies from experimentally accessible neural data.

Several computational approaches have been developed to characterize time-varying dynamics of neural interactions: state-space models^14^ capture latent dynamical structure underlying observed activity, while sliding-window correlation^15^ and coherence^16^ methods probe temporal changes in pairwise dependencies. More recent advances make use of time-varying graphical models and dynamic network inference techniques, such as adaptive Granger causality^17,18^ and Dynamic Bayesian Networks^6,19^, which model directed information flow. Among these approaches, Dynamic Bayesian Networks (DBNs) provide a particularly powerful probabilistic framework that extends traditional graphical models to capture information flow. In our previous work, we introduced multi-timescale weighted Dynamic Bayesian Networks (MTwDBN)^6^, a method that learns weighted conditional dependencies between neural populations across multiple timescales from spiking data. Because these dependencies are Granger causal, they approximate directed information flow in neural circuits. However, MTwDBN requires an *a priori* defined time window and assumes stationarity, limiting its ability to capture neural dynamics in settings where the information flow patterns evolve continuously and unpredictably.

To address this limitation, we developed continuous-time weighted Dynamic Bayesian Networks (CTwDBN), an approach that fits non-stationary DBNs and allows conditional dependencies to evolve smoothly over time. Unlike quasi-stationary models, which enforce long epochs of static connectivity punctuated by abrupt transitions, CTwDBN flexibly accommodates both smoothly evolving and discretely transitioning network structures. We validated CTwDBN on synthetic datasets and showed that it successfully recovered both continuous and discrete dynamics of information flow from observed activity. Applying CTwDBN to laminar recordings from the ventral visual stream of macaque monkeys, we found that the structure of interlaminar dependencies followed continuous trajectories through dependency space, and in contrast with the stability of receptive fields, the fluctuation persisted for hundreds of milliseconds beyond eye movements. In resting-state data, dependencies fluctuated continuously but were constrained to low-dimensional manifolds reflecting canonical motifs of laminar cortical circuitry. Together, these findings establish CTwDBN as a flexible and powerful framework for uncovering the dynamic structure of information flow.

## RESULTS

### Conceptualizing and discovering dynamic information flow in neural systems

Information flow in the brain is dynamic and shifts in response to the internal state and behavioral needs of the organism. Dynamic information flow in multivariate systems (Fig. 1a) can be conceptualized in two ways: quasi-stationary (Fig. 1b1) and continuous (Fig. 1b2). Quasi-stationarity is characterized by long periods of stability in the dependency structure of a neural network, i.e. the pattern of statistical interactions between different neural populations (Fig. 1b1). Transitions between these dependency states may or may not lead to observable changes in population firing rates, necessitating methods that can detect deeper statistical patterns to uncover state transitions. Information flow may also evolve continuously without discrete transitions, creating a smooth trajectory in a high-dimensional dependency space (Fig. 1b2). To flexibly handle both types of dynamic information flow in neural systems, we developed Continuous-Time weighted Dynamic Bayesian Network (CTwDBN), an approach that fits non-stationary DBNs and allows conditional dependencies to fluctuate smoothly in time (Fig. 1c). CTwDBN involves first using a non-stationary DBN method to calculate a family of posterior probabilities for each directed dependency in the network. To estimate this, we chose Hidden Markov Induced Dynamic Bayesian Networks^20^ (HMDBN) due to its computational speed. These posterior estimates are then summed to produce a single time-evolving probability for each directed dependency in the network. The CTwDBN can be performed in parallel over many bootstraps of the data or with different random seed initializations to improve the robustness of model fits. The output of the CTwDBN pipeline is a time-evolving network graph, where the weighted edges between nodes correspond to the probability of the directed interaction being present at a specific point in time. For continuously fluctuating systems, the network’s trajectory in dependency space can be visualized using dimensionality reduction techniques and analyzed using standard trajectory metrics. (Fig. 1d). Alternatively, for quasi-stationary systems, these graphs may be clustered to reveal discrete dependency states, with the centroids of the clusters indicating the canonical dependency structure of each state (Fig. 1e).

**Figure 1.**
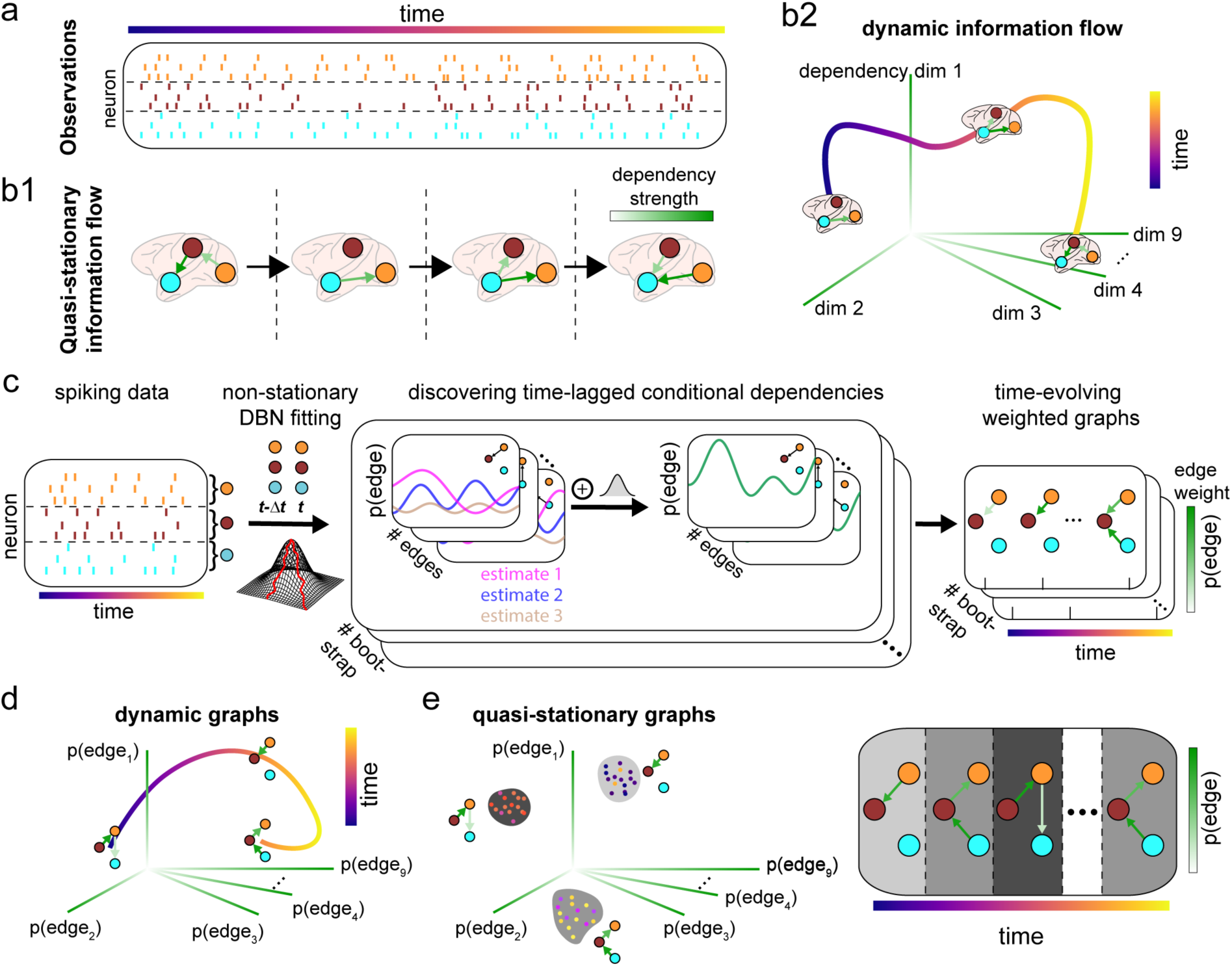
Dynamic Information Flow in Neural Systems. **(a)** Nonstationary multivariate systems can be **(b1)** quasi-stationary, i.e. exhibit periods of stable information flow with fast transitions between states. **(b2)** Alternatively, the system can move smoothly through its dependency space across time, indicating continuous, dynamic information flow. **(c)** Continuous-Time weighted Dynamic Bayesian Network (CTwDBN) pipeline for spiking neural data. A non-stationary DBN model is fit to neural data to create multiple estimates of the posterior probability of a directed interaction between two neural populations (i.e. edge). These edge probabilities are assigned as weights of the time-evolving network graphs. **(d)** For continuous dynamical systems, the trajectory of time-evolving graphs may be analyzed to reveal how neural dependencies evolve over time. **(e)** If a system transitions through discrete dependency states, time-evolving graphs will segregate into distinct clusters (left). Assigning each timepoint to a cluster reveals state transitions and the dependency structure within each state (right). Cluster timepoints are color-coded by time.

### CTwDBN recovers dynamic information flow in synthetic neural data

To validate the CTwDBN pipeline, we developed synthetic neural data that allows for the tight control of timing and information flow structure. To simulate a smoothly non-stationary system, we prepared a three-population network with connection probabilities exhibiting oscillatory dynamics over short timescales. We prepared 10,000 synthetic trials exhibiting these oscillatory dynamics and passed each trial independently through CTwDBN before averaging model outputs (see Methods). The entire network traced a figure-eight trajectory in a low-dimensional subspace that was successfully captured by CTwDBN (Fig. 2a). While the CTwDBN connection weights exhibited different scales compared to ground-truth probabilities (Fig. S1a), importantly, the shape of individual CTwDBN dependency weights matched fluctuations in ground-truth connection probabilities (Fig. 2b). Fitting the CTwDBN to a fraction of the full dataset resulted in high correlation between dependency weights and ground-truth (Fig. S1b), indicating applicability to the finite sizes of datasets regularly collected by experimentalists.

**Figure 2.**
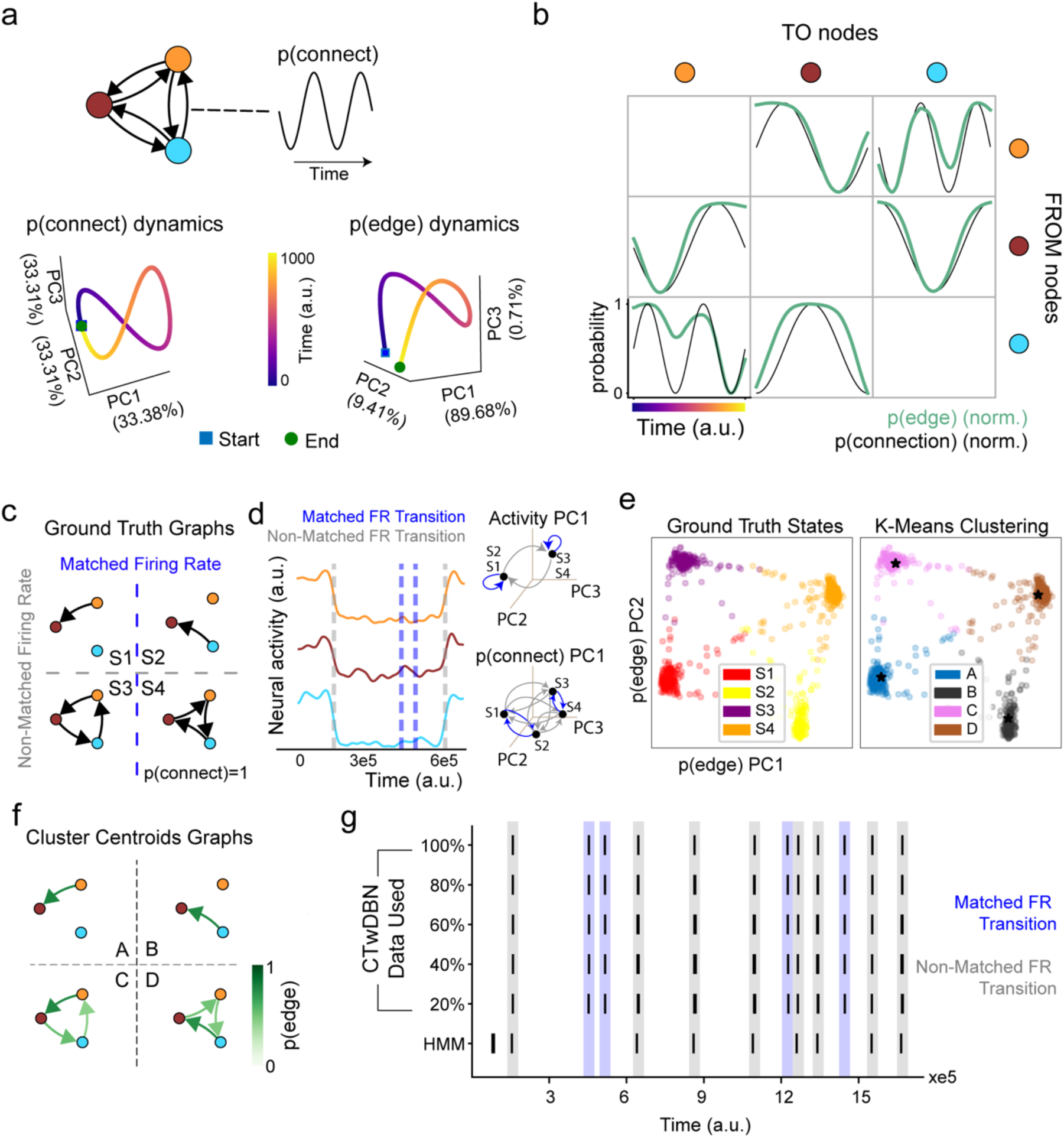
CTwDBN applied to continuous and quasi-stationary synthetic data. **(a)** A 3-population network model where the probability of interactions between nodes evolves smoothly over time (top). PCA of the model weights reveals figure-eight dynamics (bottom left) that are recovered by CTwDBN (bottom right). **(b)** Comparison of the normalized ground-truth edge probabilities (black) and the normalized CTwDBN weights (green) from the model in (**a**). **(c)** A 3-population network model with four discrete dependency states. **(d)** Neural activity from an example session snippet of a network randomly transitioning between the states in (**c**). Transitions marked by green dotted lines exhibit changes in firing rate while transitions marked by blue dotted lines do not exhibit changes to firing rate. Schematic on the right depicts the states and their possible transitions in activity space and connectivity space. **(e)** Left: PCA projection of time-evolving weighted graphs colored by ground truth state (S1-S4). Right: PCA projection of time-evolving weighted graphs colored by K-means cluster (A-D). Black stars indicate the centroids of K-means clusters. **(f)** The cluster centroid graphs identified by K-means in (**e**) match the ground truth dependency graphs in (**a**). **(g)** Raster plot of state transitions in an example session (top row) and discovered transitions by CTwDBN and a rate-based Hidden Markov Model (HMM). CTwDBN successfully discovers transitions marked by both distinct and matched firing rates, even when data is degraded through random dropping of spikes. Rate-based HMMs in contrast only detect transitions marked by changes in firing rate.

To test CTwDBN on quasi-stationary system with discrete transitions, we prepared a three-population network with four possible dependency states to transition in and out of (Fig. 2c). A subset of state transitions involved connections that were modified such that the flow of information into each population stayed constant, and there was no discernible change in the average neural activity. Other transitions involved modifications that alter the net flow of information and hence a gross change to activity levels (Fig. 2c,d). We prepared five sessions of spontaneous, long-form simulations (akin to 30 minutes at millisecond resolution) with the network randomly transitioning between these states. After fitting CTwDBN to the data, time-varying dependency graphs were clustered using K-means clustering.

Dependency graphs formed four distinct clusters confirmed by a sharp elbow in model inertia as well as a peak in silhouette score, matching the ground truth number of states (Fig. 2e, Fig. S2a,b). As visualized in two dimensions of PCA space, K-means clusters are well aligned with the ground truth state labels (Fig. 2e, Fig. S2d). We subsequently binarized the centroids of each cluster by thresholding their weighted graphs at different values, calculating the F-score as a measure of agreement with the ground-truth graph. The dependency structure of cluster centroid graphs matched the ground truth for a broad range of thresholds even when a fraction of the full data was used (Fig. 2f, Fig. S2c). In real data, where the ground truth is unknown, this threshold may be adjusted to achieve more or less conservative discovery of population connections.

Furthermore, transitions between different K-means clusters can be used to mark state transitions. In the quasi-stationary data, CTwDBN successfully identified all state transitions, regardless of discernible changes in activity levels (Fig. 2g). Furthermore, CTwDBN performance was insensitive to data degradation, with transition discovery superior to rate-based Hidden Markov Models (HMMs) even when only 20% of the available data was used to fit the model (one-sided Wilcoxon, CTwDBN > HMM across five degradations; n=5; W=15; FDR-corrected p=0.031 across all levels; Fig. S2e). This is due to rate-based HMMs only identifying state transitions marked by changes in activity levels (Fig. 2c), indicating a limitation of rate-based methods in handling states where information flow changes but activity levels remain constant.

### CTwDBN discovers dynamic inter-laminar information flow following guided saccades

To test how CTwDBN performs on real neural data, we applied CTwDBN to a dataset of laminar activity from visual area V2 of macaques performing a guided saccade task. This dataset came from prior work in our group investigating the phenomenon of receptive field remapping, the process by which the spatial regions of sensitivity (receptive fields) of visual neurons shift to their post-saccade location before the onset of eye movement. In this task, monkeys were required to execute an accurate saccade and maintain fixation on a target to obtain a juice reward. Neural receptive fields in V2 were measured before, during, and post-saccade and revealed that receptive fields across all cortical layers remap to their post-saccade location before the onset of the saccade, with remapping complete by the end of the saccade^21^ (Fig. 3a). We hypothesized that this post-saccade stability in spatial sensitivity would be reflected in the temporal dynamics of inter-laminar dependencies, and hence re-analyzed this dataset using CTwDBN for validation. Well-isolated single and multi-units in V2 were categorized by superficial/input/deep laminar identity to fit three-node CTwDBN models, and the trajectories of time-varying dependency graphs were visualized in the first three principal components (Fig. 3b).

**Figure 3.**
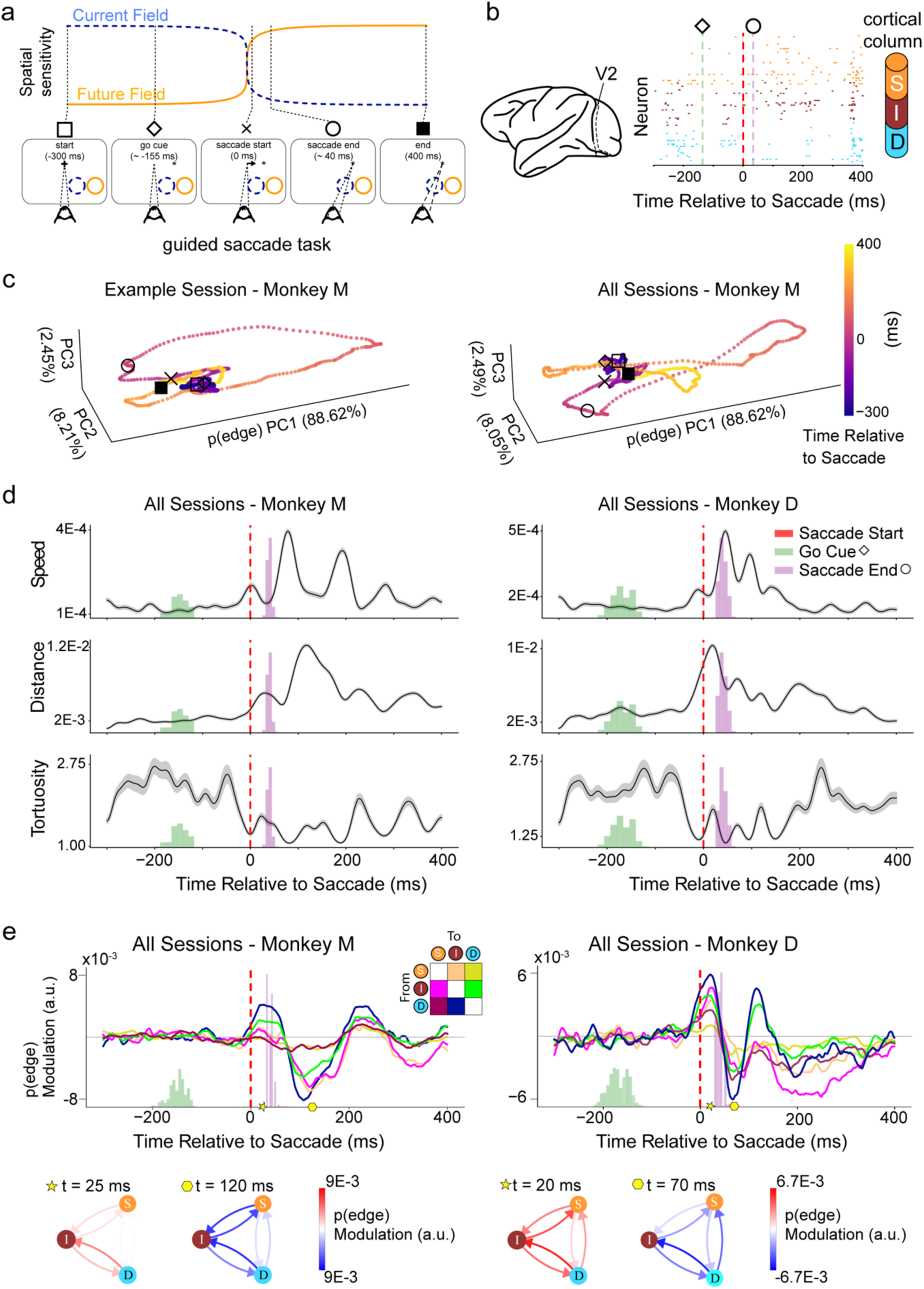
Interlaminar Dependency Trajectory Following Guided Saccades. **(a)** Changes in spatial sensitivity of V2 neurons during guided saccade task. Bottom: Task structure. The monkey maintained central fixation on a crossbar. The crossbar would disappear and a dot would appear in the periphery (random hemifield). The monkey would saccade to and maintain fixation on this dot to receive a reward. Top: Timing schematic of receptive field remapping from current (blue) to future (orange) location. Prior to initiating the saccade, V2 neurons begin to shift their receptive field by an amount equal to the saccade vector. This receptive field remapping is complete by the end of the saccade^21^. **(b)** Left: Laminar activity was recorded in visual area V2 during the guided saccade task. Neuronal populations were grouped by superficial/input/deep laminar identity to fit a three-node CTwDBN model. **(c)** Left: Dependency space trajectory of laminar populations for an example session (left) and session average (right) for Monkey M. Time zero is the beginning of saccade initiation. **(d)** Dependency space trajectory metrics of the top three PCs for the average of all sessions from monkey M (left column) and monkey D (right column). Top row: Speed is calculated as the distance traversed in PCA space divided by unit time. Middle row: Distance indicates the distance in dependency space from the beginning of the trial (t = -400 msec). Bottom row: Tortuosity quantifies the amount of twisting/winding in the trajectory. A tortuosity of one indicates a straight trajectory. Distributions for the timing of the go cue and saccade end are shown in green and purple, respectively. Red lines mark saccade initiation. Error bars show the standard error of the mean. **(e)** Inter-laminar dependency modulations relative to pre-saccade baseline (-300 - 0 msec) for monkey M (left) and monkey D (right). Error bars (standard error of the mean) are not visible due to being thinner than the mean line. Bottom: Dependency modulation graphs at key time points post saccade onset.

**Figure 4.**
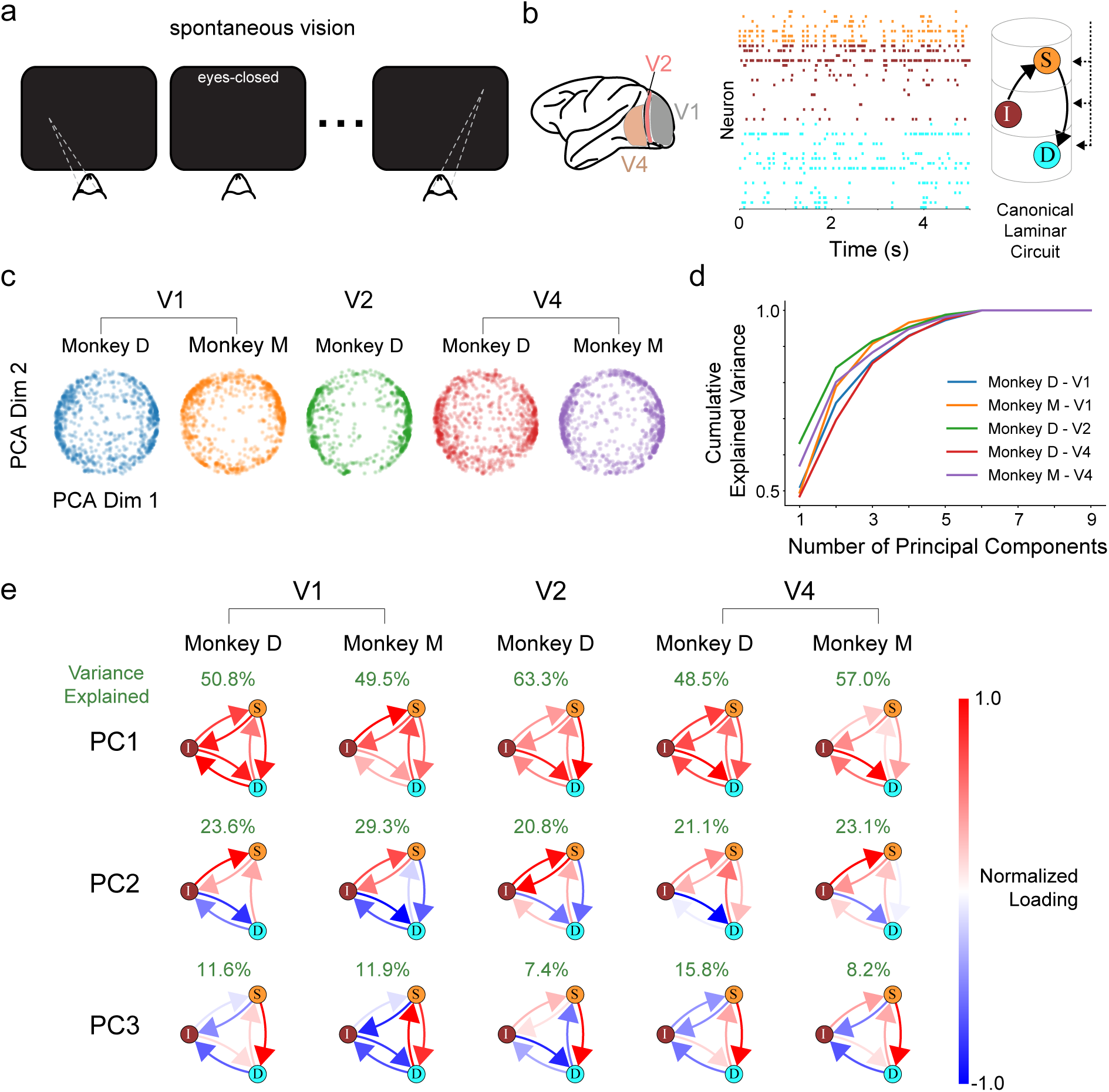
Structure of Resting Dependency Space in the Ventral Visual Stream. **(a)** Illustration of spontaneous vision. The monkey sat in a dark room and could freely move its eyes or close them for extended periods. There was no reward or imposed task structure. **(b)** Left: Laminar neural activity was recorded from visual areas V1, V2, or V4 in separate recording sessions. Right: An example segment of neural activity with neurons sorted by depth. Neurons were grouped into superficial/input/deep laminar populations to fit three-node CTwDBN models. An illustration of the canonical connectivity motifs of a cortical column with shared inputs and interlaminar connectivity is shown on the right. **(c)** PCA projection of per-timestep weighted dependency graphs for five monkey/brain region combinations. **(d)** PCA explained variance of dependency graphs in (c) for five monkey/brain region combinations. Colors match their respective condition in (c). **(e)** Graphical representation of PCA loadings for top three PCs for all monkey/brain region combinations. Percent variance explained by PC is shown above each graph.

In contrast to our findings of the stability in post-saccadic spatial sensitivity, we observed large swings in laminar dependencies following the end of saccades in both single sessions (Fig. 3c, left) and the session average level (Fig. 3c, right). To further elucidate the nature of this trajectory, we calculated the speed, distance from the start point, and tortuosity (a quantification of path twisting; lower numbers indicate straighter trajectories) over time. In both animals, we observed that the speed of the trajectory reached its peak following the end of the saccade, indicating that interlaminar connections were changing even after the eye movement had ceased (Fig. 3d, top row). The distance from the starting point (Fig. 3d, middle row) revealed that laminar dependencies deviated far from baseline following the saccade initiation and took several hundred milliseconds to return to baseline. Tortuosity was high pre-saccade and decreased during and post-saccade (Fig. 3d, bottom row), indicating that the trajectory shifted from high fluctuations to a highly coordinated, straight path in dependency space.

To interpret the large deviations in laminar dependency space identified post-saccade, we visualized the per-connection dependency modulation relative to the pre-saccade initiation period (-300 – 0 msec; Fig. 3e). This revealed several patterns consistent across both animals: a robust strengthening of laminar dependencies during the saccade and prolonged oscillations following the end of the saccade, lasting up to 400 msec after movement ceased. This contrasted with the average firing rate patterns during the same period, with neural activity decreasing before and during the saccade, followed by increased activity post-saccade (Fig. S3). This highlights a dissociation between firing-rate dynamics and inter-population dependencies. Overall, CTwDBN revealed a prolonged perturbation in inter-laminar interactions lasting hundreds of milliseconds after the execution of eye movements.

### Structure of Laminar Dependency Space in the Ventral Visual Stream of Resting Macaques

To test how CTwDBN performs on spontaneous neural data, we applied CTwDBN to a dataset of laminar activity across the ventral visual stream of macaques without any imposed task structure. Macaque monkeys sat in a dark room and were allowed to freely move their eyes or close them for extended periods (Fig. 3a). During this time, we recorded neural activity from well-isolated single or multi-units across the depth of the cortex in visual areas V1, V2, and V4 (Fig. 3b). Based on prior literature applying HMMs to neural activity of mice in unstructured environments^22,23^, we hypothesized that network graphs would cluster to reveal discrete brain states. Second, we hypothesized that information flow within these states may reflect the underlying connectivity of the laminar circuit, as resting-state functional connectivity of large brain networks has been found to reflect structural connectivity in human fMRI experiments^24^. The most prominent aspects of laminar connectivity in the sensory neocortex are common inputs to neurons across a cortical column and the canonical input → superficial → deep layer microcircuit^25,26^. Therefore, we predicted that the structure of dependency graphs would reflect these two motifs (Fig. 3b, right).

We used current-source density (see Methods) to define the boundaries of superficial, input, and deep layers, with neurons sorted accordingly to fit a three-node CTwDBN model. CTwDBN models were fit for each monkey/brain region combination separately. Since different neurons were sampled in each recording session, time-varying dependency graphs were normalized to fall onto a hypersphere (see Methods), which control analyses confirmed did not affect the clustering of synthetic data (Fig. S4).

In contrast to our first hypothesis, time-varying dependency graphs did not form discrete clusters as evidenced by the absence of a sharp elbow in K-means inertia and uniformly low silhouette scores (Fig. S5). When visualizing the first two PCs, dependency graphs tended to fall on the perimeter of a circle, indicating a low-dimensional manifold that was confirmed by the first three PCs explaining more than 75% of variance across all monkey/brain region combinations (Fig. 3d). To understand the structure of this low-dimensional dependency space in the absence of clustering, we visualized the loading of the first three PCs as graphs (Fig. 3e). Loadings refer to the contribution of each directed dependency to a principal component. Each PC represents an axis of maximum variance in the data, with loadings indicating how much each individual connection contributes to that axis. The absolute value of these loadings is arbitrary, but the relative sign of different connections reveals whether connections reflect a correlated (same sign) or anti-correlated (opposite sign) relationship between the laminar populations. All three PCs showed a striking similarity across all monkey/brain region combinations, indicating a laminar dependency space conserved across animal and cortical regions. In agreement with our second hypothesis, PCs reflected the dominant anatomical motifs of the cortical column. The first PC was characterized by a shared sign of all laminar connections, indicating that all connections increased or decreased in tandem. This general activation factor may reflect synchronization of the entire cortical column due to common inputs and high intra-columnar connectivity, and parallels the up-and down-states of firing activity identified in other state-based methods^27^. The second and third PC highlighted activation of input → superficial and superficial → deep connections, respectively, aligning with the major anatomical pathways of the canonical laminar microcircuit. These results suggest an application of CTwDBN analysis for revealing canonical structures of a network.

Taken together, these results establish CTwDBN as a general framework for characterizing how patterns of directed interactions evolve over time in neural circuits. By linking dynamic connectivity to interpretable circuit motifs, CTwDBN bridges the gap between static connectivity analyses and the temporally structured computations of neural systems. More broadly, this framework provides a foundation for quantifying how neural information flow unfolds in time, offering new leverage for studying the organizing principles of dynamic brain networks.

## DISCUSSION

The development of Continuous-Time weighted Dynamic Bayesian Networks (CTwDBN) represents a significant advancement in studying the dynamic nature of neural information flow. Unlike traditional methods that assume stationarity or quasi-stationarity, CTwDBN flexibly captures both continuous fluctuations and discrete state transitions within neural dependency structures and provides a robust and flexible framework for discovering temporal dynamics in neural systems.

Validation on synthetic data revealed that CTwDBN reliably recovers ground truth network dynamics in both continuous and quasi-stationary systems. In continuous systems, CTwDBN reliably captured the shape of the trajectory in dependency space as well as recovered dependency weights that correlated highly with ground truth connection weights, all using an amount of data that can be realistically collected experimentally. In a quasi-stationary system, CTwDBN recovered state transition times as well as the connectivity motifs of each state, even under highly degraded data conditions. This robustness is crucial in experimental studies that often contend with noise or undersampling. CTwDBN consistently outperformed traditional rate-based HMMs, highlighting sensitivity to subtle forms of neural reorganization that may be overlooked by conventional analyses.

Applying CTwDBN to experimental datasets from the macaque visual cortex provided novel insights into cortical dynamics. During guided saccade tasks, CTwDBN uncovered significant post-saccadic fluctuations in interlaminar dependencies, persisting long after the physical eye movements ceased. Interestingly, we found that neural dependencies were modulated in the opposite direction as firing rate, with an initial increase in laminar dependencies during the saccade, followed by prolonged dependency oscillations post-saccade. This may be interpreted as an increase in neural coupling during the saccade even while firing rates are suppressed. The reduction in visual cortical firing rates and visual sensitivity during eye movements are known as saccadic suppression^28^, and our findings indicate that an increase in laminar coupling may be mechanistically related to this phenomena. Spontaneous activity analysis further underscored CTwDBN’s utility by revealing a conserved low-dimensional dependency manifold across cortical regions and animals. The model fits indicated cortical circuitry operates within a constrained dynamic space, echoing established anatomical motifs. Interestingly, the lack of discrete states in spontaneous activity contrasts with previous state-based model predictions, suggesting that continuous frameworks, which do not impose state transitions, may better characterize spontaneous cortical dynamics.

### Comparison to Other Methods

CTwDBN shares conceptual similarities with recurrent switching linear dynamical systems^29^ and latent factor analysis via dynamical systems^30^, which also aim to capture non-stationary neural dynamics. However, these approaches model latent dynamics without explicitly estimating directed interactions between neural populations. Granger-based dynamic models, such as adaptive Granger causality^18^, do infer directed interactions but rely on linear autoregressive formulations and impose smooth temporal evolution of coupling through a fixed forgetting factor. Moreover, Granger causality typically estimates pairwise influences independently and does not perform joint network-level structure learning, limiting its ability to capture higher-order dependencies and abrupt network reconfigurations.

CTwDBN extends these efforts by learning coherent, interpretable, and time-resolved directed networks directly from neural data within a probabilistic graphical framework. CTwDBN accommodates nonlinear and discontinuous changes in information flow, enforces sparsity for interpretability, and quantifies uncertainty in inferred connections – all without requiring predefined state boundaries or assumptions of linearity.

### Current limitations of our approach

There are several limitations in our approach that may be improved upon in future research. As we chose HMDBN as the non-stationary DBN backbone, we inherit the Markov property where only first-order dependencies are considered, i.e. from the previous (*t-1*) to current (*t*) timepoint. We utilized a sliding window binning approach to allow for CTwDBN to discover dependencies over multiple timescales, but this prevents independent analysis of different timescales as was possible for MTwDBN^6^. CTwDBN may be extended to consider dependencies over multiple timescales by either modifying the HMDBN algorithm or switching to an alternative non-stationary method that allows for higher order dependencies such as Auto Regressive TIme Varying models (ARTIVA^31^).

### Future Directions

CTwDBN can be further generalized to incorporate behavioral data as network nodes^32^, thereby offering a unified framework for analyzing interactions between neural activity and external behavioral events. Multiple sessions of experimental data could be combined (supersessioning^33^) to study dependencies between neural populations where simultaneous recordings are unavailable, or even include synthetic variables derived from computational modeling.

Due to the generality of the CTwDBN method, any type of neural data that can be discretized is suitable for CTwDBN fitting. This includes recordings such as mesoscopic calcium imaging^34^ or fMRI that may elucidate complex network dynamics at a brain-wide scale. While model complexity increases with the number of nodes, model fitting can be simplified by incorporating priors based on known anatomy or existing theories to enhance interpretability and computational complexity.

In summary, CTwDBN provides a powerful methodological advancement for uncovering dynamic neural interactions. By explicitly capturing the fluidity of neural dependencies, CTwDBN offers a more comprehensive and interpretable view of cortical function and is flexible enough for broad application to neural systems. Continued refinement and broader application of this technique promise deeper insights into the principles underpinning cognitive flexibility, neural computation, and behavior.

## METHODS

### Continuous-Time weighted Dynamic Bayesian Network (CTwDBN) Pipeline

#### Fitting HMDBN models

The hidden-Markov–induced dynamic Bayesian network (HMDBN) algorithm was first described by Zhu & Wang^20^; readers are referred to that work for a comprehensive derivation.

##### Model formulation

HMDBN nests a Dynamic Bayesian network (DBN) inside each hidden state of a first-order hidden Markov model, allowing the directed graphical structure to change over time. Each observable variable *X_i_* is linked to a finite collection of latent “graph states” 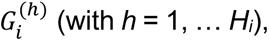 where each state specifies the *h-th* distinct parent set (i.e. the set of possible combinations of nodes interacting with *X_i_*) for *X_i_*. Transitions between states follow a node-specific Markov matrix *Ai*, and the observed data are generated from the conditional distribution table defined by the active DBN network. In this way the method relaxes the stationarity assumption of classical DBNs while preserving their causal interpretability.

##### Structure-learning strategy

Parameters and graph topology are inferred via a node-wise structural expectation–maximization (SEM) routine:

1. E-step (soft assignment). A Baum–Welch recursion estimates posterior probabilities *P*(*q_i_* (*t*) = *h* ∣ *x*) for every graph state of every node, so that each observation contributes fractionally to every candidate network epoch.
2. M-step (parameter update). Given these posteriors, closed-form re-estimates are obtained for the initial distribution *π_i_,* the transition matrix *A_i_* and the state-specific conditional tables 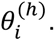
3. Structure search (inner M-step). A greedy hill-climb adds, deletes or reverses a single edge in the stationary backbone DBN, then lifts that proposal to the non-stationary space by re-identifying graph states and transition times.

Model selection uses the Baum–Welch Bayesian information criterion (BWBIC), a scoring criterion that penalizes both the number of edges and the number of hidden graphs. By weighting each sample according to its state posterior, BWBIC limits over-fitting in short segments while retaining the computational efficiency of BIC.

To robustly search the parameter space, an HMDBN model was run *B* separate times for a given dataset with a randomized starting graph (simulated data) *or* a randomized starting graph plus a different neuron sub-sample (electrophysiological data).

#### Calculation of Time-Evolving Network Graphs

For each target variable *X_j_*, HMDBN yields a catalog of candidate parent sets 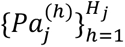 together with their time-resolved posterior probabilities *p*(*q_j_*(*t*) = *h*). To obtain a continuous measure of connection strength we defined the *marginal edge probability*

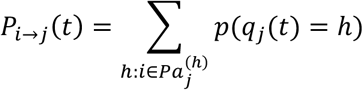

i.e. we summed the posterior mass of every graph state in which a given parent *X_i_* is present. This contrasts with the original HMDBN procedure, which located transition times by detecting crossings between competing graph-state posteriors; our marginalization avoids an arbitrary crossing threshold and preserves probabilistic information at all time points. The resulting *B* time-evolving network graphs were averaged to produce a single time-evolving network graph. This graph was optionally filtered using a zero-phase Gaussian filter implemented by FFT convolution along the temporal axis to focus on a timescale of interest. The time-evolving network graphs were stored as *N* x *N* x *T* arrays *A,* where *N* is the number of nodes, *T* is the number of time points, and the edge weight *A(i, j, t)* represents the probability of an edge from node *i* to node *j* at time *t*.

#### Principal Component Analyses (PCA)

We applied PCA (scikit-learn) to *A_vec_* for cluster visualization, analysis of PCA loadings, and trajectory analysis. All trajectory analyses were performed on the first three principal components, which are referred to here as *A_PC_*.

##### Loading Visualization

We reshaped the *N^2^* length PCA loadings to *N* x *N* and divided them by the maximum absolute value loading to normalize them between -1 and 1. We visualized the resulting *N* x *N* arrays as graphs. Because the sign of PCA loadings is arbitrary, the sign of all PCA loadings for some conditions was flipped to highlight patterns of similarity across conditions.

##### Trajectory Metrics

The instantaneous speed *S(t)* was calculated as the

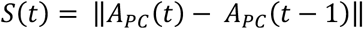

where the ||•|| denotes the L2 norm (np.linalg.norm). The distance from initial position was calculated as

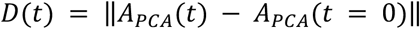

The tortuosity σ(t) was calculated on sliding windows with 1-msec steps as

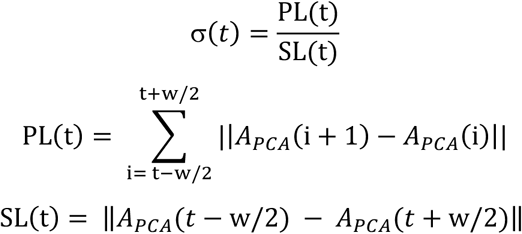

where *w* is the width of the sliding window (set to 25 msec for our analysis), *PL(t)* is the total path length traversed over *w*, and *SL(t)* is the straight-line distance traversed over *w*. To avoid division by small numbers, σ*(t)* was set to one if *SL(t)* < 1E-6.

Trajectory metrics were calculated on 100 bootstraps of the data, bootstrap size set to the total number of trials.

#### K-Means Clustering

We used K-Means clustering (scikit-learn^35^) to cluster the resulting time-evolving weighted network graphs to identify discrete states. First, graphs for each timestep in *A* were flattened to produce an *N^2^* x *T* array, *A_vec_*. To determine a suitable number of network states, we quantified the *within-cluster sum of squares* (inertia) and silhouette score for candidate solutions with K=2 … 14. For each K, we generated 100 bootstrap samples of *A_vec_* (sample size = 1000) and fitted a K-means model to each. Mean and standard error of the mean for inertia and silhouette scores were then plotted against K; the optimal K was selected at the elbow, i.e. the value of K for which the rate of decrease in inertia starts to visually slow down.

With K fixed, a single K-means run on the full time series produced

- Cluster centroids: *N^2^* vectors that represent *canonical dependency graphs*. Reshaping each centroid back to *N* x *N* yields an archetypal directed network whose edge weights can be visualized directly and optionally thresholded.
- Time-point state assignments: a label sequence *c(t)* that tags every timestep with its most probable canonical graph. Consecutive runs of identical labels demarcate stable network states, while label transitions mark putative state changes.

### CTwDBN Validation

#### A. Discrete Three-Node Network

##### Data Generation

We generated synthetic multi-population neural activity data to simulate state-dependent dynamics across interconnected excitatory-inhibitory networks. The model consisted of six neural populations organized into three modules, each containing one excitatory and one inhibitory population. Activity was simulated at 1 msec temporal resolution for 30-minute sessions.

The spike count for each population *j* at time *t* was sampled from:

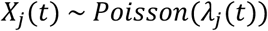

where the instantaneous firing rate *λ_j_*(*t*) was determined by:

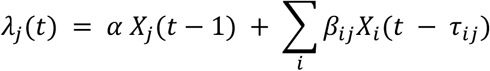

with

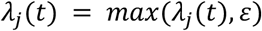

Here, *X_j_*(*t*) represents the spike count of population *j* at time *t*, *α* = 0.1 is the autoregressive coefficient (identical for all populations), *β_ij_* represents the connection strength from population *i* to population *j*, *τ_ij_* denotes the synaptic delay from population *i* to *j*, and *ε* = 0.01 ensures non-zero baseline activity.

##### Network Architecture and State-Dependent Connectivity

The system operated in one of four distinct states, each characterized by specific connectivity patterns. Within-module excitatory-inhibitory connections were preserved across all states with fixed parameters: excitatory-to-inhibitory connections (*β_ij_* = 0.5, *τ_ij_* = 2 msec) and inhibitory-to-excitatory connections (*β_ij_* = -0.5, *τ_ij_* = 2 msec). Between-module excitatory connections varied by state: State 0 featured a unidirectional connection from module 1 to module 2; State 1 featured a unidirectional connection from module 3 to module 2; State 2 implemented a feedforward chain from modules 1→2→3→1; and State 3 implemented a reverse chain from modules 3→2→1→3. All between-module connections had connection strengths *β_ij_* = 0.5 and delays *τ_ij_* = 3 msec. State transitions occurred stochastically, with each state persisting for a duration uniformly sampled between 30 and 300 seconds. Transitions were constrained to prevent immediate returns to the previous state. Initial population activity was drawn from a Poisson distribution with rate parameter *λ* = 0.1. Five independent sessions were generated.

##### Preprocessing for CTwDBN

Excitatory and inhibitory populations within each module were combined by summing their spike counts, reducing the six populations to three functional modules. Temporal binning was applied using a sliding window approach with a 3-msec window and 1-msec step size. At each time point, spike counts were summed over the preceding 3-msec window, with zero-padding applied at the boundaries to maintain the original temporal resolution of the output array. To assess the effect of sub-sampling neural data on CTwDBN outputs, *d*% (*d* ∈ {0, 20, 40, 60, 80}) of spikes were dropped from the binned data. Data from all five 30-minute sessions were concatenated to create a single super-session for each *d*. The resulting data were used to fit HMDBN models (*B* = 50 starting points).

##### CTwDBN Post-Processing

Marginal edge probabilities were filtered using a zero-phase Gaussian filter implemented by FFT (sigma = 7500 msec) convolution along the temporal axis.

##### Analysis: Comparison to Rate-Based Hidden Markov Model (HMM)

As a baseline comparison to CTwDBN, we applied traditional Hidden Markov Models to the synthetic data. We utilized Poisson HMMs implemented using the hmmlearn Python library. Similar to the CTwDBN preprocessing, excitatory and inhibitory populations within each module were combined, resulting in three variables.

For HMM analysis, spike counts were binned using non-overlapping 5-second windows. We performed extensive model selection by testing HMMs with 1 to 11 hidden states. For each number of states and each session, we fit 200 models with different random initializations. For each fitted model, we computed the log-likelihood score and Bayesian Information Criterion (BIC). The model with the highest log-likelihood score across all random initializations was selected as the best model for each session and state number combination.

##### Analysis: Calculation of Transition F-Scores

To quantify the accuracy of state transition detection, we computed F-scores between the predicted and ground-truth transition times. State transitions were identified as time points where the cluster assignment changed between consecutive samples. To avoid edge effects, transitions occurring within 15 seconds of session boundaries were excluded from analysis (corresponding to half the minimum state duration of 30 seconds). This ensured that detected transitions represented genuine state changes rather than artifacts from session concatenation.

F-scores were calculated using a temporal tolerance window of 5 seconds. A predicted transition was counted as a true positive if it occurred within 5 seconds of any ground truth transition. Predicted transitions without a matching ground truth transition within this window were classified as false positives. Similarly, ground truth transitions lacking a corresponding predicted transition within 5 seconds were counted as false negatives.

The F-Score is defined as: 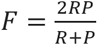 with 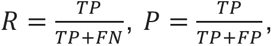 and *TP* = # model transitions which are in the ground truth, *FN* = # transitions in the ground truth not inferred by the model, *FP* = # transitions inferred by the model which are not in the ground truth. *R* and *P* refer to Recall and Precision.

##### Analysis: Calculation of Graph Centroid F-Scores

To evaluate how well K-means clustering recovered the ground-truth connectivity patterns, we computed F-scores between cluster centroids and the four known state-dependent connectivity matrices. We performed K-means clustering (k=4) on the vectorized time-evolving graphs. To assess variability and obtain robust estimates, we employed a temporal bootstrap procedure with 100 iterations. In each iteration, time points were sampled with replacement to create a bootstrap dataset of equal size to the original, and K-means was applied to identify four cluster centroids. Since K-means produces continuous-valued centroids while ground truth DAGs are binary, we applied an edge weight threshold sweep from 0.05 to 1.0 (0.05 spacing) to binarize the centroid matrices. For each threshold, centroid values exceeding the edge weight threshold were set to 1, and others to 0.

F-scores were calculated between each binarized centroid and each ground truth DAG, excluding self-connections. F-Scores were calculated using the same equation as for transition F-Scores with *TP =* # connections inferred by the model which are in the ground truth, *FN* = # connections in ground truth not inferred by the model, *FP* = # connections inferred by the model which are not in the ground truth.

To account for the arbitrary cluster labeling produced by K-means, we used the Hungarian algorithm to find the optimal one-to-one matching between clusters and ground truth states that maximized the average F-score. The mean F-score across the four matched pairs was recorded for each and bootstrap iteration. This procedure was repeated for each spike-dropping condition to assess clustering performance under varying data quality.

#### B. Continuous Three-Node Network

##### Data Generation

To evaluate CTwDBN performance on networks with smoothly varying connectivity rather than discrete states, we generated synthetic data from a 3-node binary network with time-varying edge probabilities. This model featured a continuous evolution of network structure driven by three orthogonal sinusoidal components, which endowed connections with an oscillatory structure over time.

The underlying edge probabilities evolved according to a 3-dimensional driver vector: *v(t)* = [sin(ωt), cos(ωt), sin(2ωt)]^T^, where *ω* = 2π/T ms⁻¹ where the period *T* = 1000 msec. Each of the six directed edges (excluding self-connections) was assigned a unique loading vector in this 3D space. Specifically, the forward connections from nodes 1→2, 2→3, and 3→1 were loaded exclusively on the x, y, and z components, respectively, while the reverse connections were loaded on the negative of these components. This design ensured that three principal components were required to fully characterize the network dynamics.

Edge probabilities at each time point were computed as

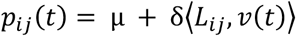

where *μ = 0.5* is the baseline probability, *δ* = 0.35 is amplitude, ⟨•⟩ represents the dot product, and *L_ij_* represents the loading vector for edge *i*→*j*. The resulting probability was clipped to [0, 1]. Self-connection probabilities remained fixed at 1.0. To avoid boundary artifacts, buffer regions of 1000 msec at the beginning and end of each trial inherited the edge probabilities from the nearest non-buffer time point.

Binary observations were generated using a logistic autoregressive model. At each time step, the probability of activation for node j was determined by:

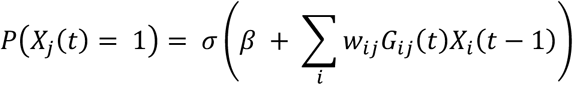

where σ is the logistic sigmoid function, *G_ij_(t)* is a binary variable indicating whether edge *i*→*j* exists at time t (sampled from the edge probabilities), *β* = -2.0 is a bias term, and edge weight *i*→*j w_ij_* = -1 if *i* == *j* and 2 otherwise. We generated 10,000 independent trials, each consisting of 3,000 time points (3 seconds at 1 msec resolution).

##### Preprocessing for CTwDBN

The binary activity data from the continuously evolving network simulations were processed at their original 1 ms temporal resolution. Each of the 10,000 trials consisted of 3 nodes observed over 3000 time points. Each trial was independently used to fit HMDBN models (*B* = 50 starting points).

##### Analysis: Sensitivity to Trial Count

To determine the number of trials required for reliable edge weight estimation, we performed a bootstrap analysis comparing CTwDBN-recovered connectivity to ground truth. CTwDBN outputs from all trials were first averaged across all starting points, yielding one connectivity matrix per trial per time point.

We focused on the central time window (1000-2000 ms) where all sinusoidal components were active to avoid edge effects, extracting weights for the six directed edges (excluding self-connections). For 1000 bootstraps, n trials (*n* ∈ {100, 250, 500, 750, 1000, 2500, 5000, 7500, 10000}) were sampled with replacement, and edge weights were averaged across the sampled trials.

Pearson correlation coefficients were computed between the trial-averaged edge weights and ground truth weights across the 1000 time points. This produced one correlation value per edge per bootstrap iteration.

#### C. Electrophysiological Recordings

Recordings were performed using 64-channel linear electrode arrays (NeuroNexus Technologies) configured with dual shanks (32 channels per shank, inter-site spacing: 70 μm, inter-shank distance: 200 μm). Prior to each experiment, electrode sites were coated with PEDOT:PSS using electroplating (nanoZ, White Matter LLC).

For each recording session, arrays were positioned in cortical area V1, V2, or V4 using an electronic micromanipulator (Narishige Inc.) under visual guidance via surgical microscopy (Leica Microsystems). This allowed accurate placement and real-time observation during insertion. The electrode traversed superficial layers at high velocity (>100 μm/s), followed by advancement at 2 μm/s through deeper tissue. After full insertion, arrays were partially withdrawn at 2 μm/s to minimize tissue compression while maintaining stable positioning.

Neural signals were acquired at 30 kHz sampling rate through a 64-channel headstage connected to an RHD Recording System (Intan Technologies). Spike sorting was performed offline using Kilosort2 with parameters: threshold = [10, 4], lambda = 10, AUC split criterion = 0.9. Automated sorting results were manually refined to identify single- and multi-unit activity.

Laminar boundaries were determined through current source density (CSD) analysis. During fixation periods, high-contrast annular visual stimuli (100% contrast) were presented for 32 ms durations, centered on receptive field locations. CSD was computed as the second spatial derivative of local field potentials and subsequently smoothed using Gaussian filtering (σ = 140 μm). Layer IV was identified by the presence of an early current sink, indicative of feedforward thalamocortical or corticocortical input. Recording sites were then classified based on their position relative to this landmark: channels located above the current sink were designated as superficial layers, while those below were classified as deep layers.

#### C1. Spontaneous Macaque

##### Data Behavioral Task

Monkeys sat in a dark room for 10 – 40 minutes (mean = 27.5 minutes, std = 8.4 min). All monitors and lights were turned off in the recording room and adjacent control room to ensure that the environment was as dark as experimentally feasible. Luminance inside the recording room was less than 9×10^−4^ cd/m^2^ (SpectraScan PR 701S, Photo Research). Eye tracker and electrophysiological data were recorded while the animals (monkeys D and M) executed eye-movements or closed their eyes freely in the dark.

##### Data Selection

Only sessions with greater than five well isolated single-/multi-units in all cortical layers (superficial/input/deep) were used for further analysis. For Monkey D this resulted in four sessions from V1 (350 units), three sessions from V2 (208 units), and seven sessions from V4 (544 units). For Monkey M this resulted in three sessions from V1 (392 units) and three sessions from V4 (207 units).

##### Preprocessing for CTwDBN

Neurons were first grouped by laminar position (superficial, input, deep) based on CSD mapping results. To ensure balanced representation across layers, we randomly sampled equal numbers of neurons from each layer. The sample size was determined by the layer with the fewest recorded neurons in a session/monkey combination. This subsampling procedure was repeated 50 times with different random seeds to ensure robustness to neuron selection. Spike counts from sampled neurons were summed within each layer, creating three population time series per session. A sliding window of 15 msec with 1 msec step size was applied to these population spike counts. Data from the same monkey/brain region were concatenated to create a single super-session. HMDBN structure learning was applied to each monkey/brain region super-session (*B* = 50 subsamples).

##### CTwDBN Post-Processing

To remove inter-session differences in baseline dependencies caused by neuron sub-sampling, the temporal mean of each marginal edge probability was subtracted for each recording session before being filtered using a zero-phase Gaussian filter implemented by FFT (sigma = 7500 msec) convolution along the temporal axis. After flattening graphs to create *A_vec_*, graphs at each timestep were normalized to unit length (sklearn.preprocessing.normalize).

#### C2. Guided-Saccade

##### Data Behavioral Task

The task design and the experimental procedures are described in detail in a previous study^21^. During the task, monkeys were required to fixate on a central point for a variable delay (500–900 msec) before making a saccade to a peripheral target that appeared abruptly. The disappearance of the fixation point served as the go cue. To receive a reward, subjects had to not only execute an accurate saccade but also maintain fixation on the target for an additional 500 msec. To discourage anticipatory saccades, both the delay duration and the target location were pseudo-randomized: delay durations were drawn from an exponential distribution, and targets appeared at one of two positions located 2.8 degrees of visual angle (dva) from the fixation point. These positions were orthogonal and each oriented 45° from the axis connecting fixation and the receptive field. Trials were only considered valid if saccades initiated within 0.75 dva of the fixation point and landed within 0.75 dva of the target. Throughout the task, Gabor stimuli with one of six orientations (0°, 30°, 60°, 90°, 120°, 150°) were presented continuously at 60 Hz on a 13×13 grid. The grid was centered 4 dva from fixation along the fixation-to-receptive-field axis. On each video frame, a single oriented Gabor appeared at a single grid location. Eye movements were detected using a velocity-based thresholding algorithm. Subjects completed an average of 895 trials (range: 729–1029) per recording session. Reaction time refers to time between go cue and saccade initiation, while saccade duration refers to the time between saccade initiation and target fixation.

##### Data Selection

Only sessions with greater than five well isolated single-/multi-units in all cortical layers (superficial/input/deep) were used for further analysis. This resulted in three sessions from Monkey D (4012 trials, 326 units) and five sessions from Monkey M (6696 trials, 572 units). All sessions recorded from area V2.

##### Preprocessing for CTwDBN

Neural activity was analyzed in 1-second windows centered on saccade onset (500 ms pre- to 500 msec post-saccade). Trials where the monkey broke fixation at any point in the trial were excluded from analysis. Neurons were first grouped by laminar position (superficial, input, deep) based on CSD mapping results. To ensure balanced representation across layers, we randomly sampled equal numbers of neurons from each layer. The sample size was determined by the layer with the fewest recorded neurons in each session. This subsampling procedure was repeated 50 times with different random seeds to assess robustness to neuron selection. Spike counts from sampled neurons were summed within each layer, creating three population time series per trial. A sliding window of 15 msec with 1 msec step size was applied to these population spike counts. HMDBN structure learning was applied independently to each trial (*B* = 50 subsamples), with trials that failed to converge excluded from subsequent analyses. This resulted in 4,591 trials from Monkey M and 2,634 trials from Monkey D. For PCA analyses, network graphs *A* were averaged over trials for either a single-session or for all sessions from the same monkey.

## AUTHOR CONTRIBUTIONS

AGS and MPJ conceptualized the project. AGS designed the synthetic data. SD and MPM collected the electrophysiological data, and ASN supervised data collection. AGS designed the data analysis pipeline and performed all analyses. MPJ supervised the project. AGS, ASN, and MPJ wrote the manuscript.

## ACKNOWLEDGEMENTS

This research was supported by NIH R01 EY034605, NIH R00 EY025026, NIH R21 MH126072 and SFARI 875855 to MPJ, NARSAD Young Investigator Grant, Ziegler Foundation Grant, Yale Orthwein Scholar Funds, NIH R01 EY032555, NIH R21 MH126072 and SFARI 875855 to ASN, NSF GRFP fellowship to AGS and by NEI core grant for vision research P30 EY026878 to Yale University. We thank the veterinary and husbandry staff at Yale for excellent animal care.

## SUPPLEMENTARY FIGURES

**Figure S1.**
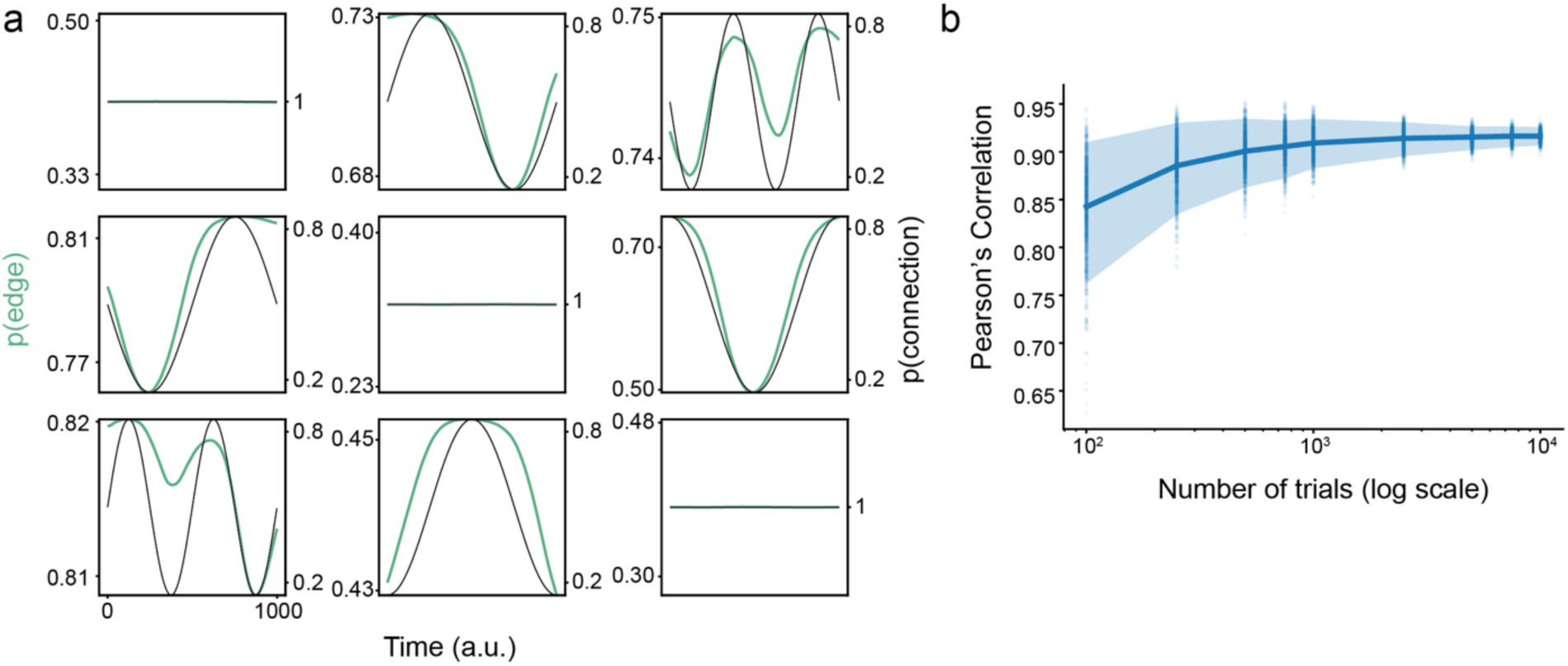
(a) Comparison of the ground-truth edge probabilities (black) and the CTwDBN weights (green). Compare to Fig. 2b. (b) Pearson’s correlation between CTwDBN weights and ground-truth edge probabilities as a function of the number of trials used to fit synthetic data. Error bar indicates 95% confidence interval.

**Figure S2.**
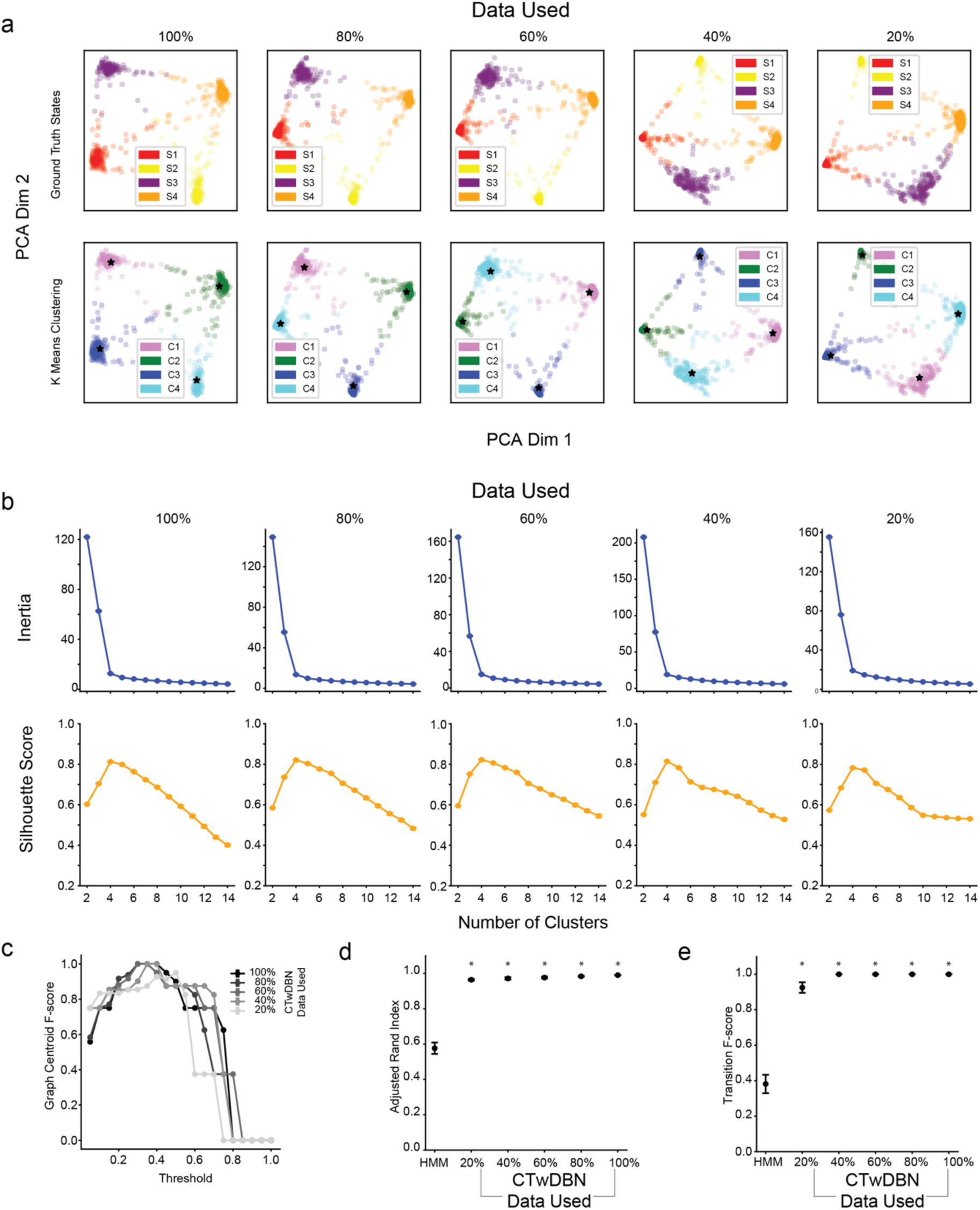
**(a)** Top Row: PCA projection of time-evolving graphs colored by ground truth state (S1-S4, top row) and by K-means cluster (C1 - C4, bottom row). Columns show CTwDBN outputs with different levels of data degradation. Black stars indicate the centroids of K-means clusters. **(b)** Model inertia (top row) and silhouette scores (bottom row) as a function of K for K-means applied to CTwDBN of varying data degradations. An optimal K of 4 is identifiable by the inertia elbow and high-magnitude peak in silhouette score at K=4. K-means performed on 100 bootstraps of the data, error bars indicate standard error of the mean. **(c)** The average F-score of graph centroids, indicating similarity between graph centroid and ground truth edge probabilities, as a function of the threshold used to binarize K-means cluster centroid graphs. Different colored lines shown for CTwDBN outputs on varying degrees of data degradation. K-means performed on 100 bootstraps of the data, error bars indicate standard error of the mean. **(d)** Adjusted rand index (indicating accuracy of assigned states) for HMM and CTwDBN outputs for differing degrees of data degradation. N = 5 synthetic sessions, error bars indicate standard error of the mean. \**p* < 0.05, CTwDBN > HMM **(e)** Transition F-score (indicating temporal alignment between predicted and actual state transitions) for HMM and CTwDBN outputs for differing degrees of data degradation. N = 5 synthetic sessions, error bars indicate standard error of the mean. \**p* < 0.05, CTwDBN > HMM

**Figure S3.**
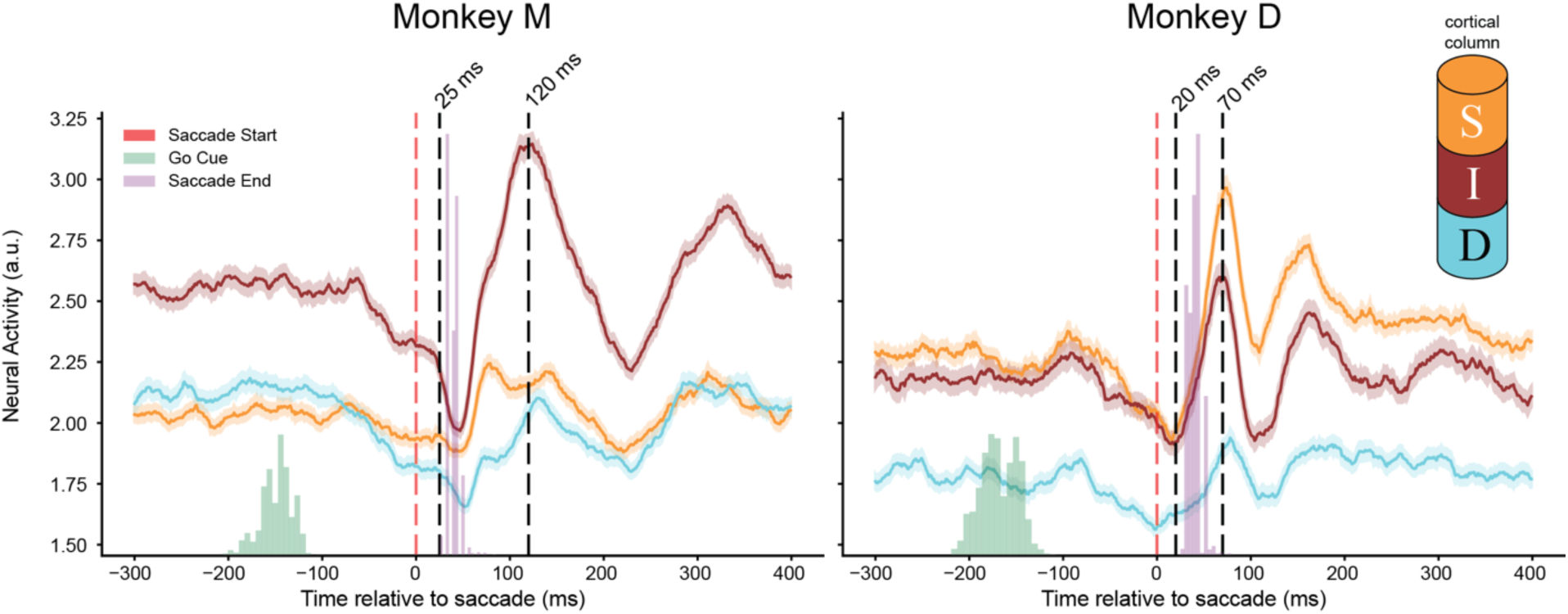
Firing rates of laminar populations during guided saccade task. Average firing rates for superficial, input, and deep populations during the guided saccade task for Monkey M (left) and Monkey D (right). Timepoints of interest (Fig. 4e) during and post-saccade are shown for each monkey separately with dotted black lines.

**Figure S4.**
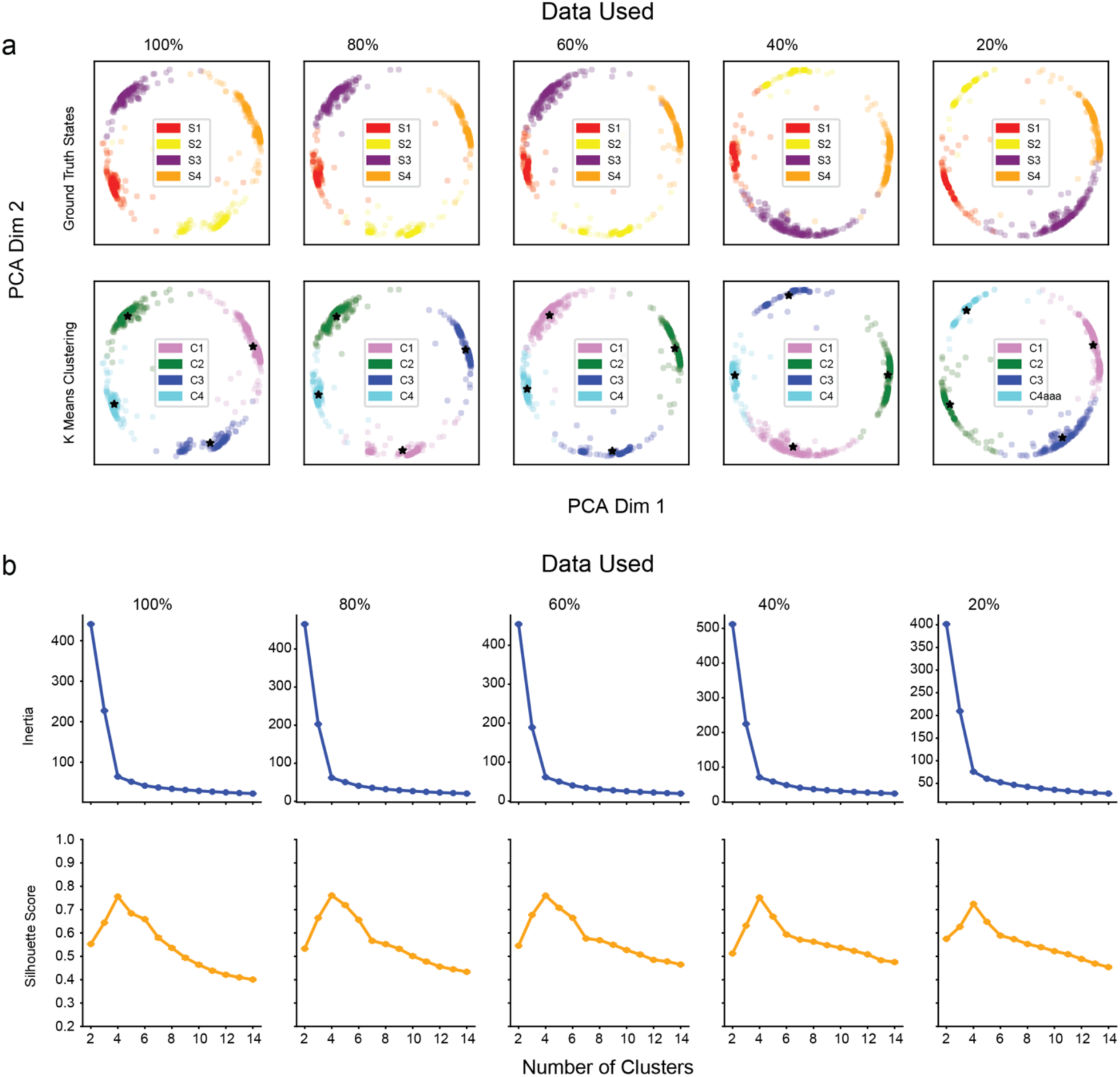
**(a)** Top Row: PCA projection of time-evolving graphs colored by ground truth state (S1-S4, top row) and colored by K-means cluster (C1 - C4, bottom row). Columns show CTwDBN outputs with different levels of data degradation. CTwDBN outputs were mean subtracted per-session and normalized to unit length vectors. Black stars indicate the centroids of K-means clusters. **(b)** Model inertia (top row) and silhouette scores (bottom row) as a function of K for K-means applied to CTwDBN of varying data degradations. CTwDBN outputs were mean subtracted per-session and normalized to unit length vectors. K-means performed on 100 bootstraps of the data, error bars indicate standard error of the mean.

**Figure S5.**
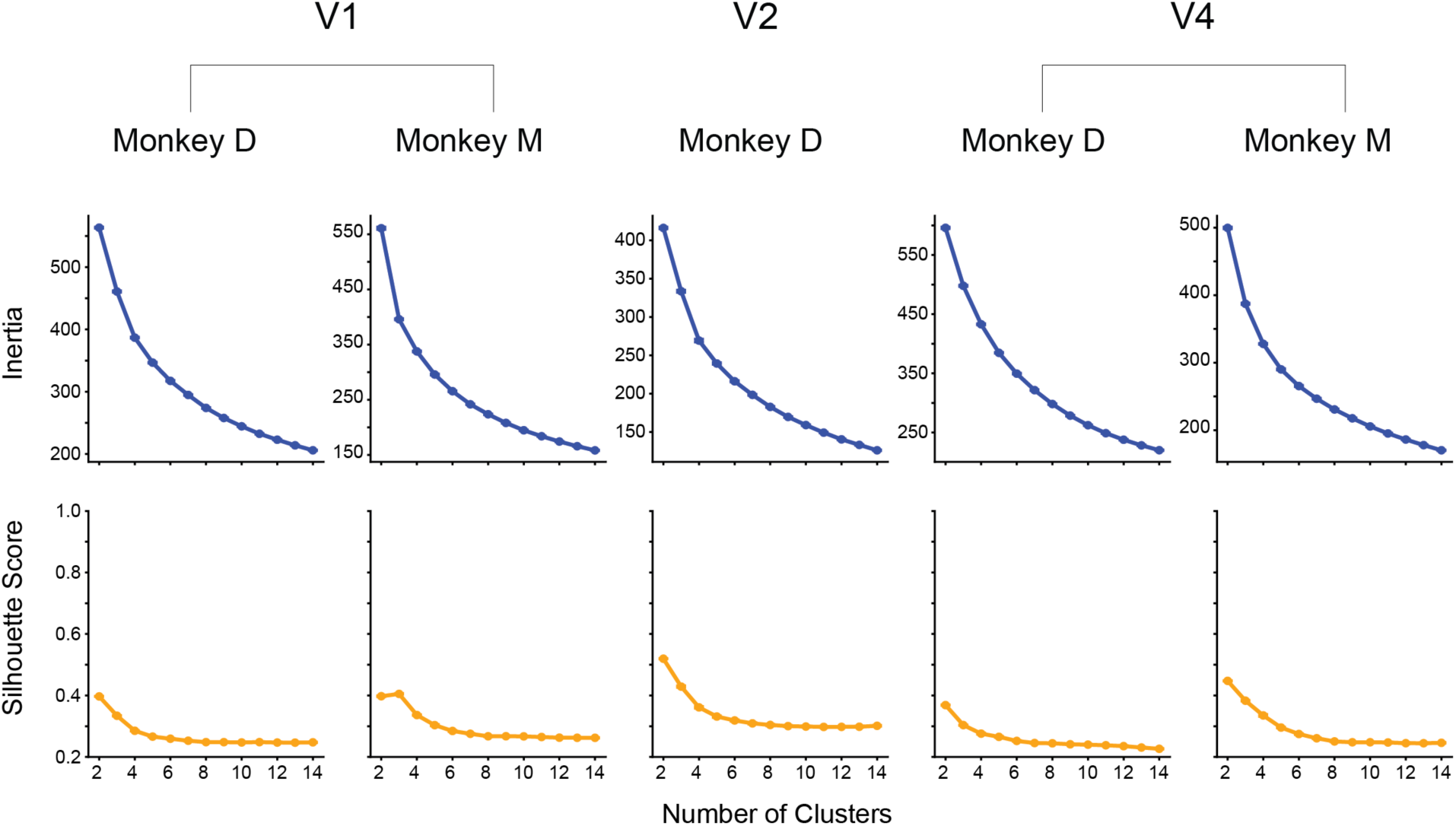
Model inertia (top row) and silhouette scores (bottom row) as a function of K for K-means applied to CTwDBN of all monkey/brain region combinations. CTwDBN outputs were mean subtracted per-session and normalized to unit length vectors. K-means performed on 100 bootstraps of the data, error bars indicate standard error of the mean.

